# CENP-C regulates centromere assembly, asymmetry and epigenetic age in Drosophila germline stem cells

**DOI:** 10.1101/2020.11.02.364794

**Authors:** Ben L Carty, Anna A Dattoli, Elaine M Dunleavy

## Abstract

Germline stem cells divide asymmetrically to produce one new daughter stem cell and one daughter cell that will subsequently undergo meiosis and differentiate to generate the mature gamete. The silent sister hypothesis proposes that in asymmetric divisions, the selective inheritance of sister chromatids carrying specific epigenetic marks between stem and daughter cells impacts cell fate. To facilitate this selective inheritance, the hypothesis specifically proposes that the centromeric region of each sister chromatid is distinct. In Drosophila germ line stem cells (GSCs), it has recently been shown that the centromeric histone CENP-A (called CID in flies) - the epigenetic determinant of centromere identity - is asymmetrically distributed between sister chromatids. In these cells, CID deposition occurs in G_2_ phase such that sister chromatids destined to end up in the stem cell harbour more CENP-A, assemble more kinetochore proteins and capture more spindle microtubules. These results suggest a potential mechanism of ‘mitotic drive’ that might bias chromosome segregation. Here we report that the inner kinetochore protein CENP-C, is required for the assembly of CID in G_2_ phase in GSCs. Moreover, CENP-C is required to maintain a normal asymmetric distribution of CID between stem and daughter cells. In addition, we find that CID is lost from centromeres in aged GSCs and that a reduction in CENP-C accelerates this loss. Finally, we show that CENP-C depletion in GSCs disrupts the balance of stem and daughter cells in the ovary, shifting GSCs toward a self-renewal tendency. Ultimately, we provide evidence that centromere assembly and maintenance via CENP-C is required to sustain asymmetric divisions in female Drosophila GSCs.

## Introduction

Stem cells are unique in that these cells can divide to give rise to daughter cells of different fates. Stem cells can undergo two distinct mitotic division types; symmetric cell division (SCD) in which the stem cell self-renews, and asymmetric cell division (ACD) in which the stem cell produces one daughter cell that undergoes differentiation [1,2]. Misregulation of the balance between SD and ACD can lead to diseases, such as cancer and infertility [3–5]. Stem cell divisions are regulated by cell extrinsic means, with well-characterised roles for signaling pathways such as Wnt, fibroblast growth factor (FGF), bone morphogenetic (BMP) [6,7]. Additionally, epigenetic mechanisms at the level of chromatin, histones and associated modifications have been implicated in the regulation of ACD. Studies in Drosophila germ line stem cells showed that prior to ACD, parental histones H3 and H4 [8,9], as well as histone H3 phosphorylated at position threonine 3 [10], are enriched on chromosomes that end up in the future stem cell. The differential distribution of histones H3 and H4 has recently been reported also in mouse embryonic stem cells [11]. These observations are in line with the ‘silent sister’ hypothesis, which proposed that sister chromatids – each carrying distinct epigenetic marks that result in differential gene expression - are selectively inherited between stem and daughter cells [12]. Moreover, the hypothesis suggested that the centromeres of each sister chromatid would also be distinct in order to facilitate selective chromosome segregation [12].

Centromeres are the chromosomal loci that specify the site of kinetochore assembly and microtubule attachment, playing a critical role in orchestrating chromosome segregation in cell division [13,14]. This locus is epigenetically defined by the incorporation of the histone H3 variant CENP-A, which is both necessary and sufficient for centromere specification and function [15–17]. Each cell cycle, newly synthesized CENP-A is assembled at centromeres to ensure functional centromere maintenance [18,19]. Recently, it has been shown in both Drosophila male and female germ line stem cells (GSCs) that CID assembly shows unique properties [20,21]. Firstly, CID is deposited at centromeres prior chromosome segregation, during G_2_/prophase, a cell cycle time that is distinct compared to symmetrically diving cells [20,21]. Secondly, CID is unevenly distributed between sister centromeres, with between 1.2-1.5 fold more CID inherited by ‘stem’ side sister chromatids [20,21]. A third line of evidence showed that parental CID – as opposed to newly synthesized CID - is found to be enriched in both intestinal and germline stem cells [21,22]. Finally, studies in GSCs showed that the mitotic spindle is asymmetric both temporally and with respect to the distribution of microtubules; at prometaphase sister chromatids of the future stem cell attach first to the spindle and more spindle microtubule are observed in the stem cell side at metaphase [20,21]. Taken together, these studies propose a model by which CID asymmetry can drive the selective attachment of microtubules leading to the non-random segregation of sister chromatids [23,24].

Further investigations into how the mitotic chromosome segregation machinery -and specifically centromeres - are altered in asymmetric divisions are now needed. Indeed, relatively little is known about centromere assembly and maintenance in stem cells. In addition to CID, the Drosophila centromeric core is comprised of two key components, the inner kinetochore protein CENP-C and the centromere assembly factor CAL1 [25,26]. CAL1 binds to CID-H4 dimers and assembles CID nucleosomes [27–29]. CENP-C binds to CID containing nucleosomes, and also interacts directly with CAL1, recruiting new CAL1-CID-H4 to the centromere [28–30]. In addition, CAL1 can then recruit new CENP-C to the centromere, closing the epigenetic loop [28,29]. In Drosophila GSCs, both CAL1 and CENP-C are asymmetrically distributed between stem and daughter cells [20,21]. Functional experiments – either overexpression or depletion - have shown that CAL1 is required to maintain CID asymmetry in GSCs, impacting on cell fate and development [20,21]. CENP-C is also critical for the assembly and maintenance of CID/CENP-A at fly and human centromeres [25,31,32]. Yet, whether CENP-C can regulate stem cell asymmetric division beyond its canonical mitotic kinetochore function remains unclear. In this study, we investigate CENP-C function in Drosophila GSCs. We find that CENP-C is required for CID assembly in GSCs, as well as maintaining appropriate CID asymmetry between stem and daughter cells. In addition, we determine CID and CENP-C levels to decrease in accordance with GSC age. We propose that CENP-C’s function in CID assembly and asymmetry maintains the balance of symmetric and asymmetric divisions in the GSC niche impacting on long term GSC maintenance in the ovary.

## Results

### CENP-C is assembled at GSC centromeres in G_2_/prophase

At the apical end of the Drosophila germarium (Fig 1A), 2-3 GSCs are found attached to cap cells (Fig 1B). Female GSCs divide asymmetrically to give a differentiating daughter cell called a cystoblast (CB) [33]. We previously showed that CID is assembled at GSC centromeres between the end of DNA replication up until at least prophase [20]. To assess the cell cycle timing of CENP-C assembly in GSCs, we used 5-ethynyl-2′-deoxyuridine (EdU) incorporation to mark cells in and out of S-phase and 1B1 staining to mark the spectrosome, the shape of which can be used to define the cell cycle stage [34,35] (Fig 1C, 1D). We focused on GSCs with a pan nuclear EdU staining pattern characteristic of mid to late S-phase, in which the spectrosome forms a bridge shape (Fig 1C). GSCs that were EdU negative with a round spectrosome and with centromeres distributed throughout the nucleus, but without condensed chromosomes, were deemed to be in G_2_/prophase (Fig 1D). We then quantified total CENP-C fluorescent intensity (integrated density) at centromeres at both stages (Fig 1E). We found an increase in CENP-C level between cells in S-phase, compared to cells in G_2_/prophase. Quantitation revealed an average increase of 38% in CENP-C (S-phase=23.36±1.84, n=32 cells; G_2_/prophase=32.14±1.611, n=34 cells). These results indicate that similar to CID, CENP-C is assembled at GSCs centromeres between the end of S-phase and G_2_/prophase.

**Figure 1:**
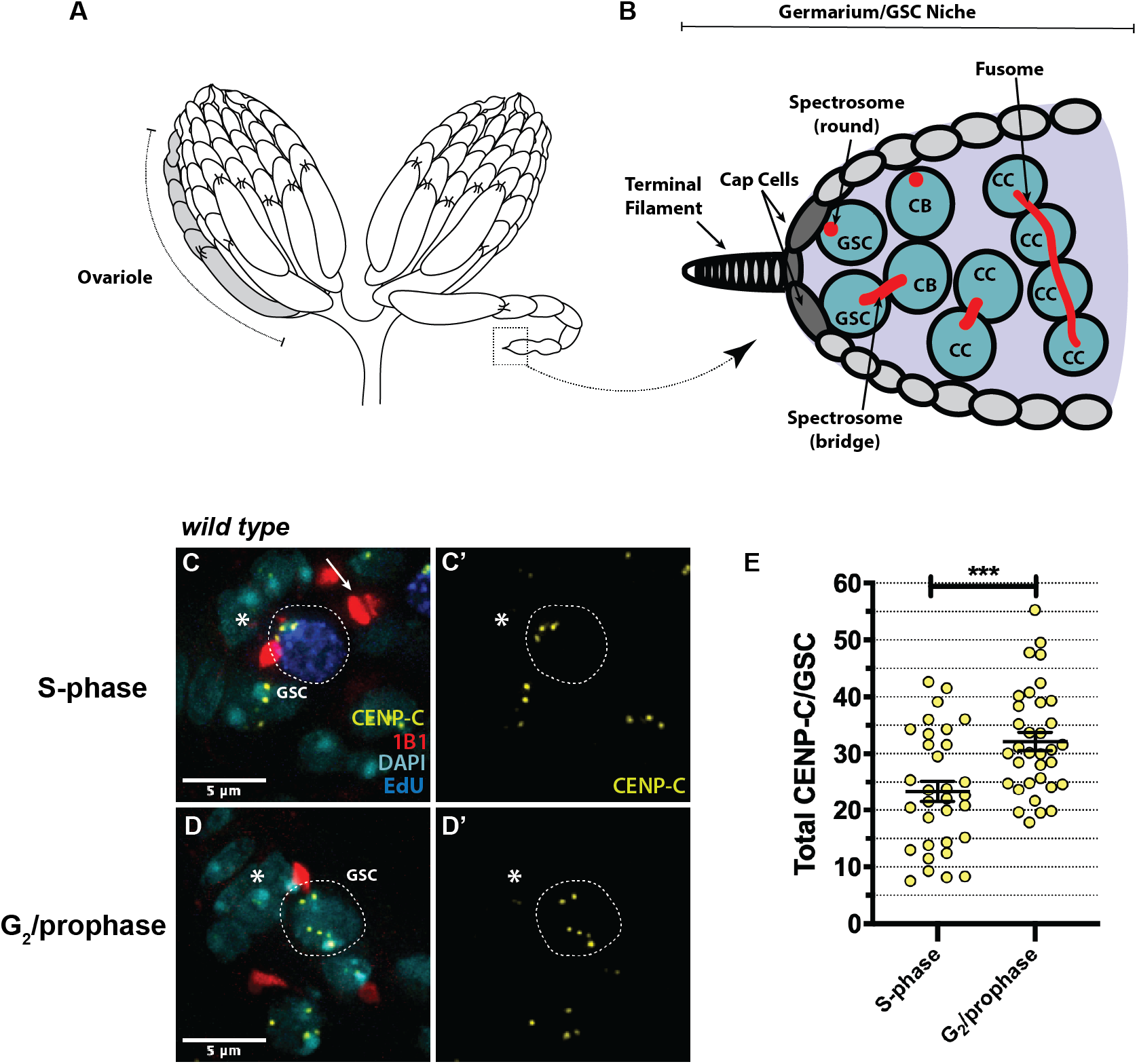
CENP-C is assembled between S-phase and G_2_/prophase in female GSCs. (A) Schematic of the Drosophila ovary, composed of 16 ovarioles (ovariole highlighted in grey) organised into developing egg chambers. The GSC niche is located in the anterior-most chamber of the ovariole, the germarium (boxed). (B) Schematic of the GSC niche and 2- and 4-cell cysts in the germarium. G_2_/prophase GSCs can be identified with a round spectrosome attached to the cap cells. CB= cystoblast, CC= cystocyte. (C-D) Immunofluorescent image of wild type GSCs (circled) in S-phase (C, C’) and G_2_/prophase (D, D’) stained with DAPI (cyan), EdU (blue), spectrosome (1B1, red) and CENP-C (yellow). Arrow represents spectrosome associated with the circled S-phase GSC. (E) Quantitation of total CENP-C fluorescent intensity (integrated density) in GSCs at S-phase and G_2_/prophase. ***p<0.001. Scale bar = 5 μm. Error bars = Standard Error of the Mean (SEM).

### CENP-C is required for CID assembly in the germline, specifically in GSCs

To determine if CENP-C is required for CID assembly in GSCs, we used the GAL4-UAS system to induce the RNAi-mediated depletion of CENP-C using the germline-specific driver *nanos-GAL4*. To confirm CENP-C knock down, control *nanos-GAL4* and CENP-C RNAi ovaries were stained with antibodies against CENP-C and 1B1 to mark the spectrosome (Fig S1A-D’). We quantified the total CENP-C fluorescent intensity (integrated density) in GSCs with a round spectrosome, indicative of cells in G_2_/prophase (Fig S1E). Quantitation revealed an approximate 60% depletion of CENP-C in GSCs (*nanos-GAL4*=34.45±1.65, n=29 cells; CENP-C RNAi=13.70±1.63, n=28 germaria). We next labelled and quantified CID fluorescent intensity in GSCs depleted for CENP-C, both at S-phase and at G_2_/prophase (Fig 2A-D). As expected in the *nanos-GAL4* control, CID intensity increased between S-phase and G_2_/prophase (S-phase=15.82±0.73, n=40 cells; G_2_/prophase=24.58±1.45, n=43 cells), by approximately 35% on average (Fig 2I). However, in the CENP-C RNAi we did not observe this increase (S-phase=17.46±1.06, n=36 cells; G_2_/prophase=15.50±0.96, n=43 cells) (Fig 2I). Indeed, CID levels were comparable between S-phase and G_2_/prophase. This result indicates that CENP-C is specifically required for CID assembly that occurs between the end of S-phase up to prophase in GSCs.

**Figure 2:**
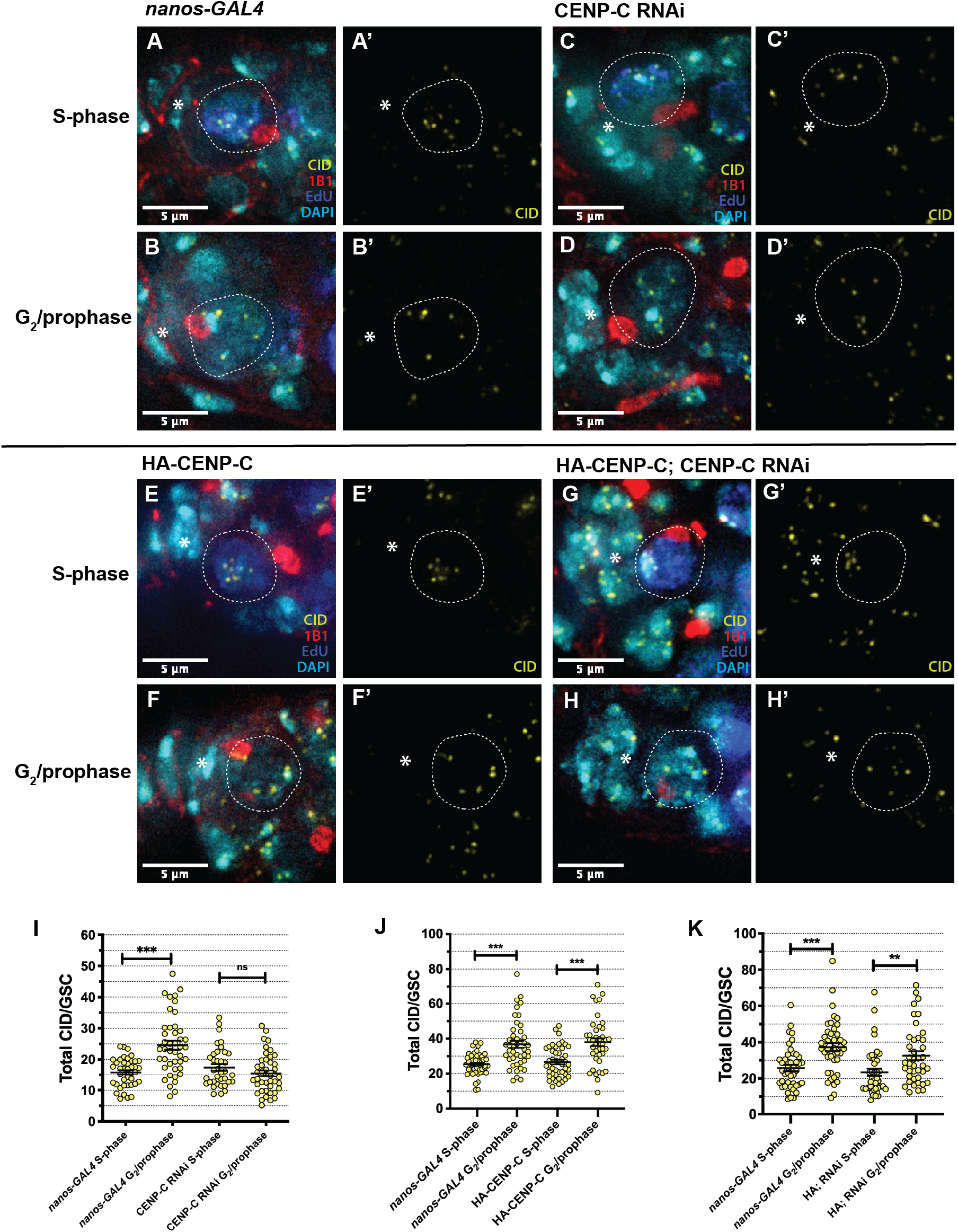
CENP-C is required for CID assembly in GSCs. (A, B) *nanos-GAL4*, (C, D) CENP-C RNAi, (E, F) HA-CENP-C and (G, H) HA-CENP-C; CENP-C RNAi (rescue) stained with DAPI (cyan), EdU (blue), 1B1 (red) and CID (yellow). S-phase GSCs (A, C, E, G) are positive for EdU, contain a bridge spectrosome and clustered centromeres. G_2_/prophase GSCs (B, D, F, H) are EdU negative, contain a large round spectrosome and dispersed centromeres. GSCs are circled. * denotes cap cells. Scale bar = 5 μm. Quantitation of total CID fluorescent intensity (integrated density) in GSCs at S-phase and G_2_/prophase in *nanos-GAL4* and (I) CENP-C RNAi, (J) HA-CENP-C and (K) HA-CENP-C; CENP-C RNAi. ***p<0.001, **p<0.01, ns=non-significant. Error bars = SEM.

We next investigated whether the localisation of the CID assembly factor CAL1 was affected by CENP-C knockdown. For this, we antibody stained control and CENP-C-depleted germaria for CAL1, as well as CENP-C in order to distinguish centromeric from nucleolar CAL1 (Fig S2A-C’). Both centromeric and nucleolar CAL1 was visible in the *nanos-GAL4* and CENP-C RNAi. However, we quantified total centromeric CAL1 in GSCs at G_2_/prophase and found centromeric CAL1 to be reduced in the CENP-C RNAi compared to the *nanos-GAL4* control (*nanos-GAL4*=16.01±1.45, n=30 cells; CENP-C RNAi=9.44±0.85, n=29 germaria) (Fig S2D). This result is in line with the structural evidence that CENP-C is the recruitment factor for CAL1-CID-H4 complexes, marking the centromere for new CID assembly [28]. Lastly, to assess if CENP-C is required for CID localisation at later stages of development, we knocked down CENP-C using the *bam-GAL4* driver active in 4-8 cell cysts. As previously reported [20], staining for CENP-C revealed an effective knock down at this stage (Fig S2E-H’). Surprisingly, we noted that at this stage knockdown of CENP-C did not lead to a major reduction in CID level, nor any noticeable phenotype (Fig S2I-L’). This finding is comparable to our previous observations for CID and CAL1 [20], and suggests that CENP-C function in centromere assembly might be dispensable for later divisions occurring the germarium.

### Excess CENP-C does not promote additional CID assembly in GSCs

To further monitor CENP-C function in CID assembly in GSCs, we overexpressed an N-terminal HA-tagged CENP-C using the *nanos-GAL4* driver (Fig S1F-I). To measure the extent of CENP-C over-expression, we labelled total CENP-C (tagged and endogenous) with an anti-CENP-C antibody and quantified its intensity in *nanos-GAL4* and HA-CENP-C GSCs at G_2_/prophase (Fig S1J). Here, total centromeric CENP-C level increased by approximately 45% (*nanos-GAL4* =31.86±2.59, n=22 cells; HA-CENP-C=51.02±4.49 n=20 cells). We then measured CID assembly between S-phase and G_2_/prophase in the background of increased CENP-C (Fig 2E, 2F). In GSCs overexpressing HA-CENP-C, CID intensity increased at the expected rate between S-phase and G_2_/prophase, in line with the *nanos-GAL4* driver (*nanos-GAL4_S-phase_*=25.44±0.88, n=48 cells; *nanos-GAL4_G2/prophase_*=36.77±2.01, n=45 cells; HA-CENP-C_S-phase_=26.63±1.28, n=46 cells; HA-CENP-C_G2/prophase_=37.99±2.20, n=41 cells) (Fig 2J). Moreover, fluorescence values between control and HA-CENP-C are comparable, indicating that increased CENP-C level does not correlate with increased CID assembly in GSCs. We next designed rescue experiments, in which we overexpressed HA-CENP-C that is resistant to the shRNA in the CENP-C RNAi background (Fig 2G, 2H, S1A’’-D’’). Upon over-expression, we quantified total CENP-C levels, comparing the HA-CENP-C; CENP-C RNAi to that of *nanos-GAL4* and CENP-C RNAi (Fig S1E). Here, ‘rescued’ GSCs displayed an 85% restoration of total CENP-C levels (*nanos-GAL4*=34.45±1.65, n=29 cells; CENP-C RNAi=13.70±1.63, n=28 germaria; HA-CENP-C; CENP-C RNAi=29.49±2.09, n=28 germaria). Finally, we measured CID assembly between S-phase and G_2_/prophase in the HA-CENP-C; CENP-C RNAi background (Fig 2K). In this case, CID assembly was partially rescued, displaying an increase in CID level from S-phase to G_2_/prophase, although not quite to the CID level in the control (*nanos-GAL4_S-phase_*=25.68±1.76, n=43 cells; *nanos-GAL4_G2/prophase_*=37.24±1.98, n=44 cells; HA-CENP-C;CENP-C RNAi_S-phase_=23.37±1.98, n=42 cells; HA-CENP-C;CENP-C RNAi_G2/prophase_=32.51±2.47, n=41 cells). These results show that over-expression of CENP-C alone does not affect CID assembly, but CENP-C expression rescues the defect in CID assembly observed in the CENP-C RNAi.

### CENP-C functions to maintain an asymmetric CID distribution between GSCs and CBs

Our previous characterisation of centromere positioning in GSCs and CBs at anaphase and DNA replication, allowed us to conclude that both cells enter synchronously into S-phase immediately at the end of mitosis, without a detectable G_1_ phase [20]. We also showed that in addition to CID, CENP-C is asymmetrically distributed between GSC-CB S-phase ‘pairs’ [20]. Here, we again confirmed this result for CID in *nanos-GAL4* (Fig 3A), and next tested if CENP-C is required for the asymmetric distribution of CID (Fig 3B). For this, we measured CID intensity in GSC-CB S-phase pairs, expressed as a ratio of total CID in GSC/CB, in CENP-C-depleted GSCs compared to the control *nanos-GAL4* (Fig 3A, 3B, S3A). Quantitation revealed a significant increase in the GSC/CB ratio of CID intensity to 1.44 in the CENP-C RNAi versus 1.2 in controls (*nanos-GAL4_GSC/CB_*=1.19±0.06, n=40 cells; CENP-C RNAi_*GSC/CB*_=1.44±0.08, n=36 cells (Fig 3E). This indicates that CENP-C functions not only in CID assembly in G_2/_prophase, but also in the maintenance of CID asymmetry in S-phase. Given that new CID assembly is likely deficient in the absence of CENP-C (Fig 2), this result indicates a potential bias in the retention of parental CID by the GSC. We next investigated CID asymmetry upon HA-CENP-C overexpression (Fig 3C). Comparing the ratio of total CID in GSC-CB pairs in S-phase, quantitation showed no significant change asymmetry (Fig 3F, S3B). In this case, HA-CENP-C overexpression did not significantly affect the GSC/CB ratio (*nanos-Gal4*_GSC/CB_=1.22±0.06, n=47 cells; HA-CENP-C_GSC/CB_=1.12±0.04, n=46 cells) (Fig 3F). To verify that the shift in CID asymmetry to 1.44 was dependent on CENP-C, we performed the same analysis in the HA-CENP-C; CENP-C RNAi rescue line (Fig 3D). Indeed, quantitation of rescue versus *nanos-GAL4* controls returned the expected CID ratio of 1.2 (*nanos-Gal4*_GSC/CB_=1.23±0.07, n=43 cells; HA-CENP-C; CENP-C RNAi_GSC/CB_=1.20±0.06, n=42 cells) (Fig 3G, S3C). These results show that CENP-C functions to maintain the correct level of CID asymmetry in stem and daughter cells at S-phase.

**Figure 3:**
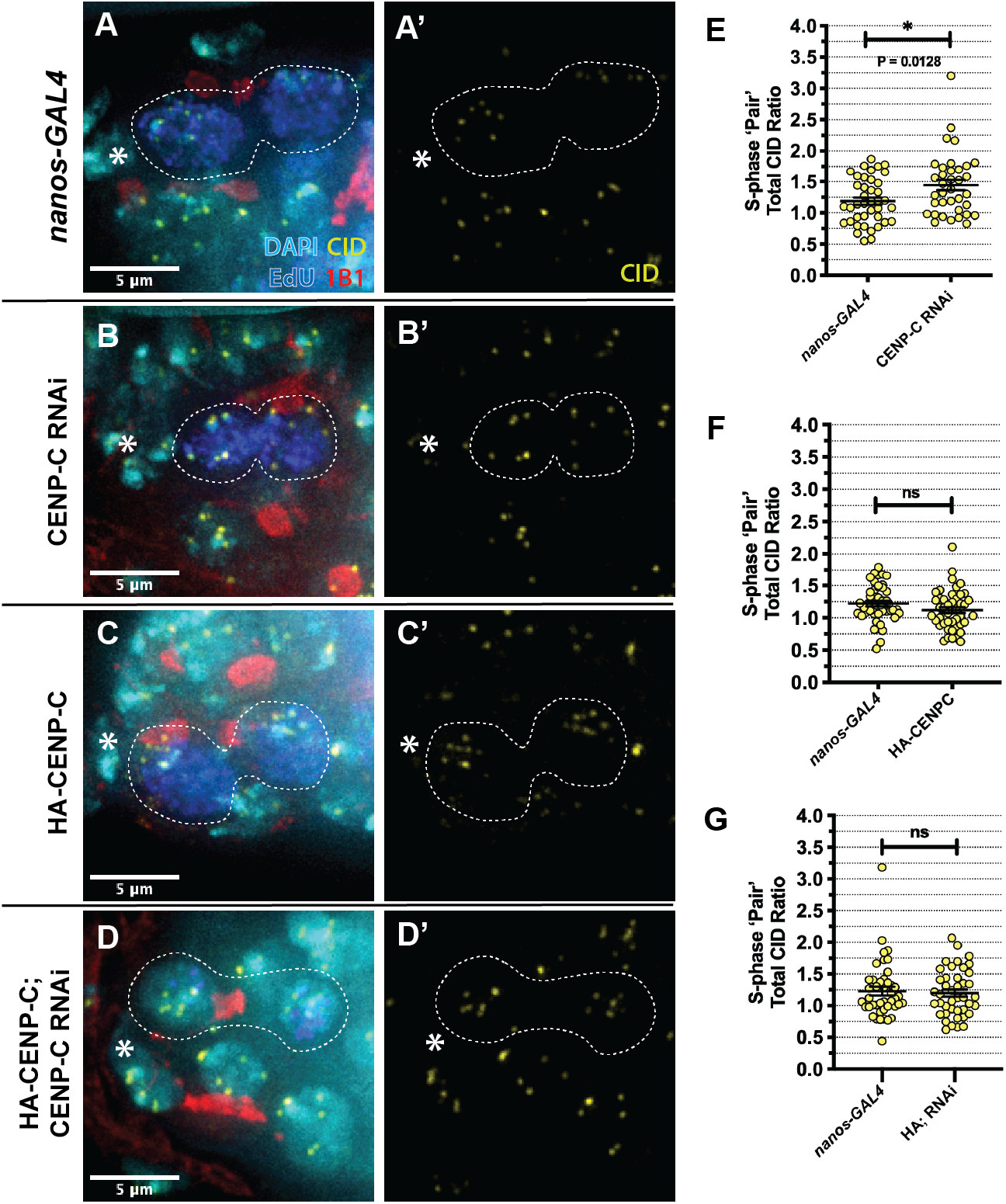
CENP-C is required for asymmetric CID distribution between GSCs and CBs at S-phase. (A) *nanos-GAL4*, (B) CENP-C RNAi, (C) HA-CENP-C and (D) HA-CENP-C; CENP-C RNAi (rescue) stained with DAPI (cyan), EdU (blue), 1B1 (red) and CID (yellow). S-phase GSCs and CBs are positive for EdU, contain a bridge spectrosome and clustered centromeres. Dashed white line outlines GSC/CB pairs. * denotes cap cells. Scale bar = 5 μm. Quantitation of the ratio of total CID fluorescent intensity (integrated density) between GSC/CB S-phase pairs in *nanos-GAL4* and (E) CENP-C RNAi, (F) HA-CENP-C and (G) HA-CENP-C; CENP-C RNAi (rescue). Each point represents the ratio of total CID between GSC versus its corresponding CB. ns=non-significant. *p<0.05. Error bars = SEM.

### CENP-C regulates GSC proliferation and long term GSC maintenance

To probe the function of CENP-C in GSCs proliferation or maintenance, control and CENP-C knockdown ovaries were stained for the germ cell marker VASA, as well as 1B1 marking round spectrosomes and branched fusomes. Control *nanos-GAL4* germaria contained the expected GSCs and germ cell content, in line with previous studies [20]. In contrast, CENP-C depleted germaria revealed a spectrum of germ cell proliferation phenotypes (Fig 4A-D). Quantitation of phenotypes (Fig 4E) showed that over one third of germaria (35%) analysed 5 days after eclosion showed normal development, comparable to the control. However, another third (32%) displayed an accumulation of germ cells, indicative of germ line tumours [36]. The final third (29%) displayed isolated GSC and CBs located in the niche and 4-8 cell cyst stages were lacking, indicative of a differentiation defect. Finally, a small proportion of germaria (4%) lacked GSCs entirely. Analysis of germaria 10 days after eclosion revealed an exacerbation of the GSC loss phenotype (21%) (Fig 4E). Importantly, HA-CENP-C overexpression almost completely rescued the differentiation defect and GSC loss phenotypes at 5 days, when expressed in conjunction with the CENP-C shRNA (Fig 4E). These results suggest that CENP-C is required for GSC proliferation, as well as the long-term maintenance of the GSC population.

**Figure 4:**
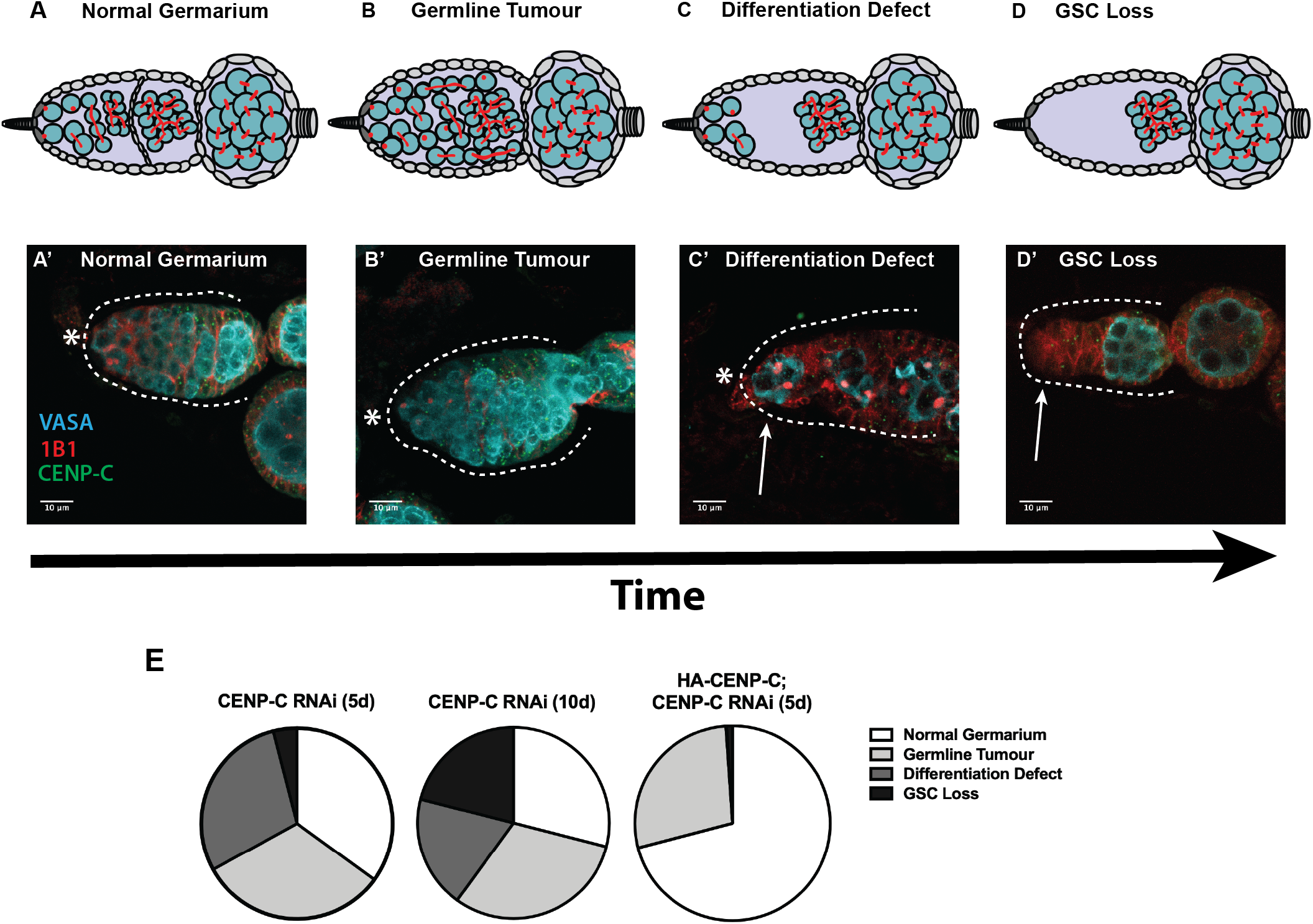
CENP-C depletion disrupts GSC proliferation and maintenance over time. (A-D) Characterisation of the phenotypes arising in CENP-C depleted germaria. (A, A’) Normal germarium are healthy with the expected lineage of germ cysts and spectrosome/fusome development. (B, B’) Germline tumours are characterised by an increased number of germ cells (GSCs, CBs, cysts) in the germaria, often displaced from their normal position with abnormal spectrosome/fusome morphology. (C, C’) The differentiation defect is characterised by a pool of GSCs/CBs in the apical end of the germaria (white arrow), separated from later stage developing cysts. (D, D’) GSC loss is characterised by the absence of GSCs (and often CBs and early germ cysts; white arrow) at the apical end of the germarium. * denotes cap cells. Scale bar = 10 μm. (E) Quantitation of the frequency of the above phenotypes observed at 5-days (5d) and 10-days (10d) post-eclosion, and in the HA-CENP-C; CENP-C RNAi rescue at 5-days (5d) post-eclosion. Charts each represent 3 biological replicates (50 germaria analysed per replicate).

We next investigated if the accumulation of germ cells after CENP-C depletion might be due to a cell cycle block. Given that CENP-C normally functions in kinetochore attachment to microtubules, we assayed whether cells were blocked in mitosis using phosphorylation at serine 10 of histone H3 (H3S10P) to marks cells at late G_2_-phase to metaphase. CENP-C depleted germaria displaying the GSC tumour phenotype were generally negative for H3S10P (Fig S4A-H). H3S10P combined with DAPI staining of DNA allowed us to monitor for chromosome segregation defects, which were also absent (data not shown). These results show that the extent of CENP-C depletion (60% reduction) does not result in a mitotic arrest nor in chromosome segregation defects, indicating that the canonical kinetochore function of CENP-C is maintained. Moreover, mitotic delay or arrest does not explain the observed cell proliferation phenotype. We then used EdU incorporation to label cells with or without newly replicated DNA as a marker of S-phase. Strikingly, we noted that EdU positive 2-, 4- or 8-cell cysts were more frequently observed in CENP-C depleted germaria (Fig S4I-P). Quantitation showed that while 0-3 EdU positive cysts (median of 1) were observed in *nanos-Gal4* germaria, this number increased in CENP-C RNAi (median of 2) (Fig S4Q). This suggests that cysts in the CENP-C RNAi progress slower through DNA replication in S-phase perhaps contributing to the observed accumulation of germ cells in germaria.

### CID and CENP-C levels are reduced in aged GSCs and CENP-C reduction accelerates CID loss

*Wild type* GSCs retain 1.2-fold more CID in an asymmetric division. Given that symmetric GSC divisions (in which the CID ratio is presumably 1) occur at a low frequency [37,38], we hypothesised that CID and CENP-C levels would gradually be depleted in GSCs over time (Fig 5). We investigated this possibility in *wild type OregonR* GSCs dissecting at 5-, 10- and 20-days post-eclosion, staining for 1B1 to mark GSCs in G_2_/prophase and either CID (Fig 5A-C) or CENP-C (Fig 5E-G). Quantitations showed a dramatic 45% decrease in CID level between 5- and 20-day timepoints (*OregonR*_5-day_=28.73±1.72, n=28 cells; *OregonR*_10-day_=20.62±1.29, n=30 cells; *OregonR*_20-day_=16.22±0.97, n=30 cells) (Fig 5D). Similarly, CENP-C reduced at a comparable rate at each timepoint (*OregonR*_5-day_=28.22±1.35, n=28 cells; *OregonR*_10-day_=19.46±1.23, n=29 cells; *OregonR*_20-day_=15.44±1.11, n=29 cells) (Fig 5H). Hence, CID and CENP-C levels in GSCs reduce in correlation with GSC age. We next wanted to determine if this observed reduction in CID was dependent on CENP-C. For this, we quantified CID in 5- and 10-day old germaria in both *nanos-GAL4* and CENP-C RNAi GSCs at G_2_/prophase (Fig 5I-L). In the CENP-C RNAi, we quantified germaria displaying normal and germline tumour phenotypes at 5-days old and the differentiation defect phenotype at 10-days old. In *nanos-GAL4* GSCs controls, we observed a reduction in total CID signal between 5- and 10-days old (*nanos-GAL4*_5-day_=31.46±1.80, n=29 cells; *nanos-GAL4*_10-day_=24.50±1.19, n=32 cells) (Fig 5M), comparable with *wild type* observations. In CENP-C-depleted GSCs at 5- and 10-days old, we found that CID was reduced further (CENP-C RNAi_5-day_=20.21±1.40, n=33 cells; CENP-C RNAi_10- day_=14.62±1.41, n=28 cells). Moreover, the percentage decrease in CID between control 5- and 10-day old GSCs (19%) was amplified even further upon knockdown of CENP-C (28%). These results support a role for CENP-C in long-term CID maintenance in aged GSCs.

**Figure 5:**
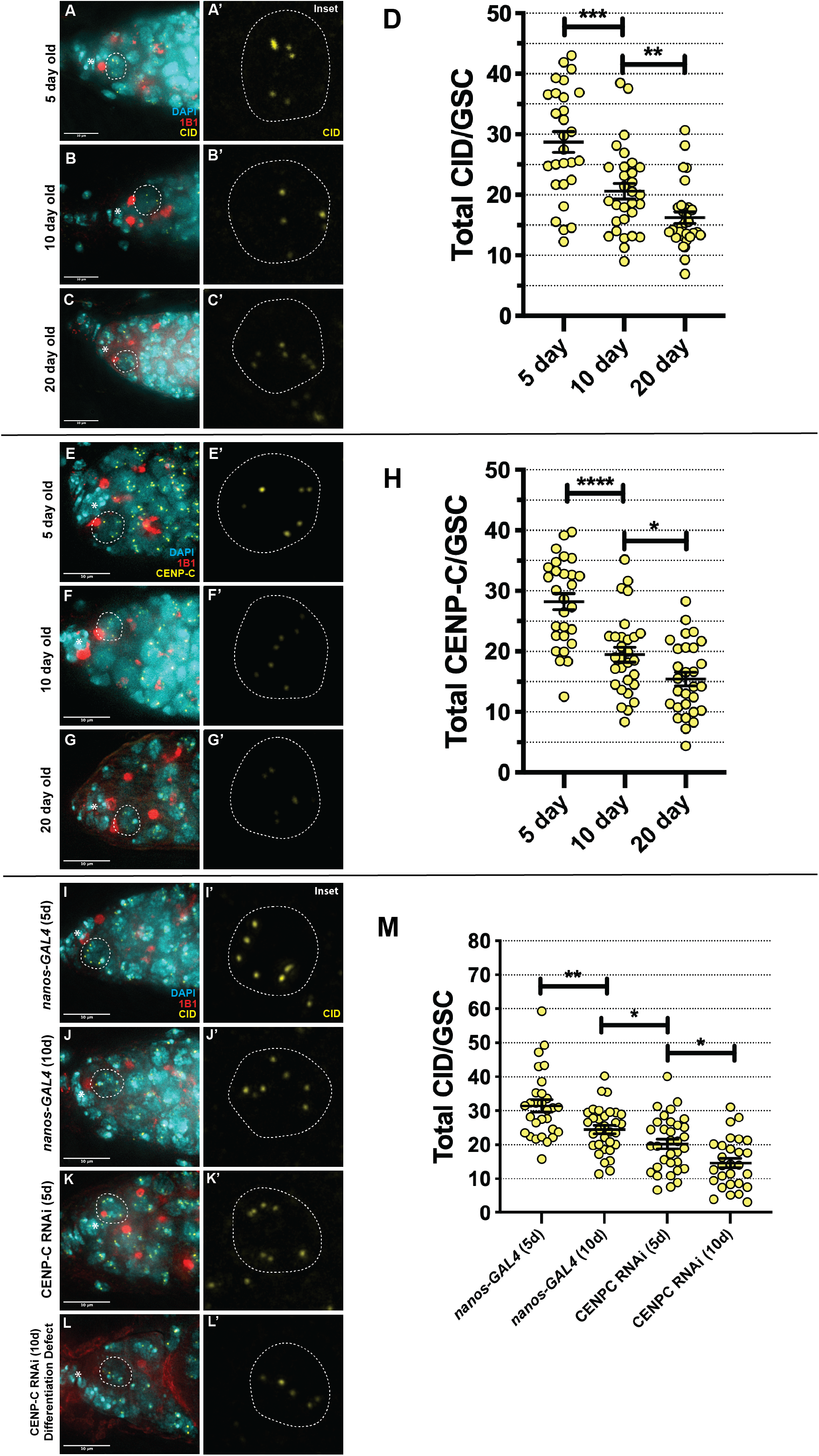
CID and CENP-C level is reduced in aged GSCs and CENP-C depletion accelerates CID loss. (A-C) *Wild type* germaria (5-, 10- and 20-day old) stained with DAPI (cyan), 1B1 (red) and CID (yellow) or (E-G) CENP-C (yellow). GSCs are boxed and inset. * denotes cap cells. Scale bar = 10 μm. (D) Quantitation of total CID (D) or CENP-C (H) integrated density in *wild type* GSCs at 5-, 10- and 20-days post eclosion. *p<0.05, **p<0.01, ***p<0.001, ****p<0.0001. Error bars = SEM. (I-L) Germaria of *nanos-GAL4* (5d, 10d), CENP-C RNAi (5d, 10d differentiation defect phenotype) stained with DAPI (cyan), 1B1 (red) and CID (yellow). GSCs are boxed and inset. * denotes cap cells. Scale bar = 10 μm. (L) Quantitation of total CID integrated density per GSC in *nanos-GAL4* (5d, 10d), CENP-C RNAi (5d, 10d differentiation defect phenotype). *p<0.05, **p<0.01. Error bars = SEM.

### CENP-C regulates the balance of GSCs and CBs in the niche

To specifically explore CENP-C function in GSC maintenance, we assayed the GSC/CB balance in CENP-C depleted germaria. To measure GSC/CB balance, we used the stem cell marker pMad [39] and SEX-LETHAL (SXL) that labels the GSC-CB transition up to the 2-cell cyst (2cc) stage [40,41] (Fig 6A-F and Fig S5A-D). Firstly, in *OregonR* (*wild-type*) and RNAi isogenic control lines, we counted the number of pMad-positive and SXL-positive cells in each germaria at 5-days (Fig S5E). We next used this data to calculate the SXL/pMad ratio as a measure for the number of GSCs compared to CBs and 2ccs in each germarium (Fig S5F). In both controls, although the number of positive pMad and SXL cells differ (Fig S5E), the SXL/pMad ratio remained similar, with approximately 4 SXL-positive cells for every 1 pMad-positive cell at 5-days old (Fig S5F). Analysis of *OregonR* germaria at 10- and 20-days old revealed an unexpected gradual decrease in the SXL/pMad ratio (*OregonR*_10day_3.55 ± 0.16; *OregonR*_20day_3.12 ± 0.12) and thus a change in the balance in stem/daughter cells over time (Fig S5F). In *nanos-GAL4* 5-day old germaria, we counted approximately 1.5 pMad-positive cells and 6 SXL-positive cells on average (Fig 6G). Therefore, *nanos-GAL4* controls have 4 SXL-positive cells for each pMad-positive cell at 5-days post eclosion (4.15 ± 0.21) (Fig 6H). At 10 days, this ratio dropped in a manner similar to *wild type* (3.69 ± 0.21) (Fig 6H). In the CENP-C RNAi germaria analysed at 5-days post-eclosion, the number of pMad-positive cells increased to 2.5 on average, while the number of SXL-positive cells did not change (Fig 6G). As a result, the SXL/pMad ratio is reduced to 2.7:1 (2.68 ± 0.17) (Fig 6H). This ratio for CENP-C RNAi is further reduced to 2.0:1 at 10-days post eclosion (1.99 ± 0.14) (Fig 6H). In contrast, overexpression of HA-CENP-C alone did not change the SXL/pMad ratio. In this case, HA-CENP-C expressing germaria dissected at 5-days old showed approximately 2 pMad positive cells on average, but approximately 8 SXL positive cells (Fig 6G). Thus, the SXL/pMad ratio remained normal at approximately 4 (3.78 ± 0.19) (Fig 6H). Significantly however, HA-CENP-C overexpression was sufficient to almost fully rescue the disrupted SXL/pMad ratio observed in the CENP-C RNAi (3.58 ± 0.12) (Fig 6H). Taken together, these results indicate that (1) the balance of stem/daughter cells in the niche slowly changes with age, and (2) GSCs with reduced CENP-C shift the balance of stem/daughter cells toward self-renewal rather than differentiation, offering an explanation for the differentiation defect we see at 5- and 10-days old (Fig 4A-E).

**Figure 6:**
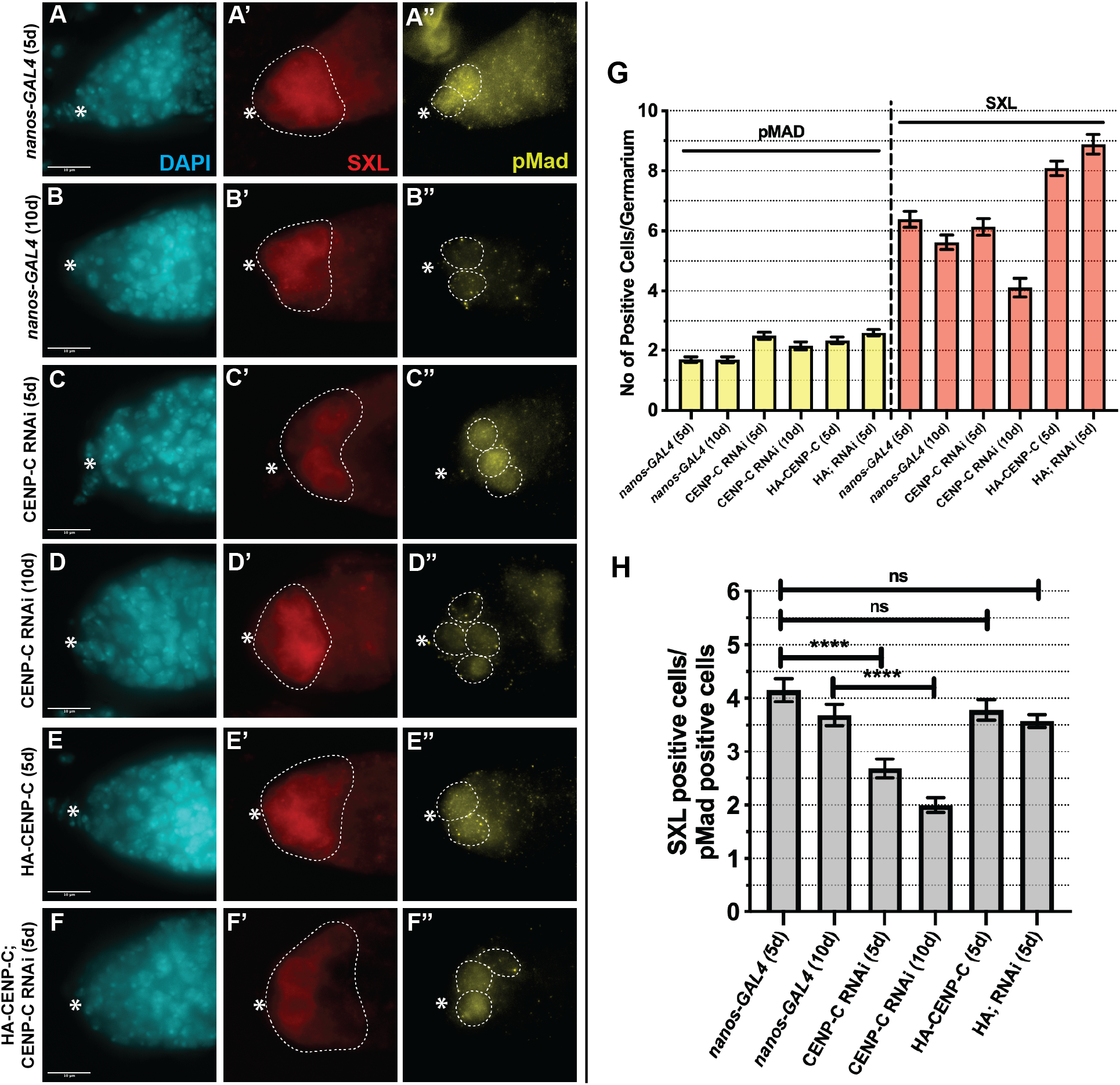
CENP-C depletion shifts GSCs toward a self-renewal tendency. (A-F) *nanos-GAL4* (5d, 10d), CENP-C RNAi (5d, 10d), HA-CENP-C (5d) and HA-CENPC;CENPC RNAi rescue (5d) germaria stained with DAPI (cyan), SEX-LETHAL (SXL, red) and pMad (yellow). Scale bar = 10 μm. * denotes cap cells. White dashed circles highlight SXL or pMad positive cells. (G) Quantitation of the number of pMad positive (yellow) and SXL positive (red) cells per germarium. (H) Ratio of the number of SXL:pMad positive cells per germarium. **** p<0.0001. ns = non-significant. Error bars = SEM.

## Discussion

### CENP-C contributes towards mitotic drive by facilitating CID assembly, maintaining CID asymmetry and assembling a strong GSC kinetochore

Drosophila GSCs use the strength differential between centromeres to bias sister chromatid segregation between stem and daughter cells [20,21]. This asymmetry in centromere strength is achieved through differential CID assembly in G_2_/prophase, which is used to build a stronger kinetochore and mitotic spindle [20,21]. We have previously shown that CENP-C is asymmetrically distributed between GSC-CB pairs after cell division [20]. We now show that similar to CID, CENP-C is also assembled in G_2_/prophase of the cell cycle. Moreover, we find that CENP-C is required for CID assembly at this cell cycle time. This direct role for CENP-C in CID (CENP-A) assembly has been previously characterised, mostly in cultured cells [25,28,42]. However, few studies have investigated aberrant centromere assembly in stem cells or in the context of tissue development. Here, we show that defective CID/CENP-C assembly has a profound effect on GSC maintenance and in turn oocyte development over time. In addition to its function in assembly, we find that CENP-C is required to maintain correct asymmetry of CID between stem and daughter cells. Specifically, depletion of CENP-C gives rise to GSCs retaining 1.44-fold more CID compared to 1.2 in the controls. Previously CAL1 overexpression (together with CID) in female GSCs resulted in a CID ratio of 1 [20]. Taken together, it appears that distorting CID asymmetry (to either 1 or 1.4) disrupts the balance of stem and daughter cells in the ovary. How CENP-C functions together with CAL1 to maintain the correct level of asymmetry remains unclear, however it may relate to different requirements for CAL1 and CENP-C in maintaining pools of newly synthesized or parental CID at distinct cell cycle times. Ultimately, our findings for CENP-C function in GSCs are in full agreement with the mitotic drive model for stem cell regulation [9].

### How might parental CID be maintained in stem cells?

Previous studies in Drosophila male GSCs and intestinal stem cells (ISCs) have shown that parental CID, as opposed to newly synthesized CID, is preferentially maintained by stem cells [21,22]. Given the lack of centromere assembly in CENP-C-depleted GSCs in our case, we can deduce that these GSCs contain largely parental CID. Furthermore, our results suggest a bias in the retention of parental CID by the stem cell. Many questions surround how parental CID could be maintained at centromeres. In the male germline, testes-derived DNA and chromatin fibres display a high frequency of unidirectional fork movement [9], providing a potential mechanism as to how old versus new histone asymmetry might be established and maintained. Given the timing for CID assembly in G_2_/prophase (after DNA replication and sister centromere establishment), parental CID is redistributed at the replication fork before new CID assembly. It is therefore likely that parental CID requires direct maintenance via CENP-C, CAL1 or other histone chaperones. Intriguingly, in CENP-C-depleted germaria, we frequently observed germ cell cysts (GSC, CB, 2cc, 4cc, 8cc) in S-phase, suggesting a potential non-canonical function for CENP-C at this cell cycle time. Indeed, previous photobleaching experiments in human cell lines showed that unique from most other centromere proteins CENP-C is stable during S-phase [43]. More recently, CENP-C has been shown to maintain centromeric CENP-A in S-phase and allow for error-correction of CENP-A assembly at non-centromere sites [44]. Furthermore, HJURP (functional CAL1 equivalent in humans) is required to maintain CENP-A during DNA replication [45]. It is tempting to speculate that CENP-C (and/or CAL1) might be utilised in S-phase to maintain parental CENP-A in an asymmetrically dividing stem cell system.

### Adult stem cells age epigenetically at the centromere

Numerous studies have shown that the epigenome changes with age (‘epigenetic drift’), particularly related to DNA methylation, histone modifications and chromatin remodeling [46,47]. Importantly, this epigenetic ‘erosion’ also pertains to stem cells [48,49]. In this context, an epigenetic regulator of aging should ideally decrease over time and directly influence cell fate. Here we show that both CID and CENP-C decrease approximately 45% on average between 5- and 20-days old in *wild type* GSCs. This loss is further exacerbated upon a reduced CENP-C level, suggesting that CENP-C is directly involved in this centromere ‘erosion’. It is likely that the low frequency of symmetric stem cell divisions [37,38] (and in turn symmetric CID distribution) gradually depletes these centromere proteins over time. To our knowledge, this is the first time that the centromere has been implicated in stem cell aging and is consistent with an early study showing centromere loss in aged women [50].

### Centromeres as regulators of stem cell fate and differentiation

By measuring the ratio of stem to daughter cells, we show firstly that the balance of stem to daughter cells in the niche changes gradually over time. Secondly, disruption to the centromeric core by depletion of CENP-C shifts the balance towards GSC self-renewal (reducing the SXL/pMad ratio), and this is exacerbated further over time. Later in development, we observe differentiation defects, measured by an absence of germ cell cysts in the germarium. Thus, CID level is closely linked with stem cell self-renewal rate, which in turn manifests in a failure in differentiation at cyst-stages, followed by GSC loss. Indeed, CENP-C has been previously implicated in Drosophila stem cell maintenance and/or differentiation, being 1 of 42 genes identified in three different RNAi screens [51–53]. Recently, it has been shown that reprogramming human fibroblasts to pluripotency results in a removal of CENP-A from the centromere [54]. Moreover, low levels of CENP-A prevent human pluripotent stem cells from differentiating, resulting in continuous self-renewal [55]. This implies a certain centromere ‘load’ required to differentiate – a prospect reinforced by our observations in the germline. Thus, we propose the role of the centromere on cell fate is two-fold, with 1) centromere ‘load’ and 2) parental CID/CENP-A pools being key regulators in stem cell fate. How CENP-A load ultimately leads to a change in gene expression should be a focus of future studies.

## Materials and Methods

### Fly stocks and husbandry

Stocks were cultured on standard cornmeal medium (NUTRI-fly) preserved with 0.5% propionic acid and 0.1% Tegosept at 20°C under a 12 hours light-dark cycle. All fly stocks used were obtained from Bloomington Stock Centre (#) unless otherwise stated. The following fly stocks were used: *Oregon-R* (#2371), *wild-type* (#36303, RNAi isogenic control), *nanos-GAL4* (#25751), *bam-GAL4* (kind gift from Margaret T. Fuller), UAS-CENP-C RNAi (#38917), UASp-HA-CENP-C;*SM6 Cy* (kind gift from Kim S. McKim), HA-CENP-C; UAS-CENP-C-RNAi (this study). CENP-C knockdown (and rescue) using the *nanos-GAL4* driver was performed at 22°C and using the *bam-Gal4* driver at 29 °C. HA-CENP-C was induced using *nanos-GAL4* at 25 °C. F_1_ progeny were dissected 5, 10 or 20 days after eclosion. Results obtained from each experiment rely on three biological replicates, unless otherwise specified.

### Immunofluorescence (IF)

After fixation, samples were immediately washed in 1XPBS-0.4% Triton-X100 (0.4% PBST). Samples were then blocked in 0.4% PBST with 1% BSA for 2-4 hours at room temperature, incubated with primary antibodies (in blocking buffer) overnight at 4 °C. Samples were then washed in 0.4% PBST for 3x 30 minutes. Secondary antibodies are added (1:500 in blocking buffer) for 2 hours at room temperature in the dark. Samples are again washed 3x 30 minutes in 0.4% PBST followed by addition of DAPI (1:1000) for 15 minutes in 1XPBS.

### EdU Incorporation

Ovaries were dissected and incubated for 30 min with EdU (0.01 mM) in 1XPBS and then fixed as described. After washing in 0.4% PBST, ovaries were incubated for 30 minutes in the dark with 2 mM CuSO_4_, 300 μM fluorescent azide and 10 mM ascorbic acid. Samples were then washed with 0.4% PBST for 10 minutes and then blocked and stained as described above.

### Antibodies

For immunostaining, the following antibodies were used: rabbit anti-CENP-A (CID) antibody (Active Motif 39719; 1:1000), sheep anti-CENP-C (Dattoli *et al*, 2020; 1:2000), mouse anti-H3S10P (Abcam ab14955; 1:1,000), rabbit anti-VASA (Santa Cruz sc-30210; 1:300), rat anti-VASA (Developmental Studies Hybridoma Bank (DSHB); 1.500), mouse anti-Hts (1B1, DSHB; 1:500), rabbit anti-CAL1 (Bade *et al*, 2014; 1:1000), rabbit anti-SMAD3/5 (pMad) (Abcam; 1:500), mouse anti-SEX-LETHAL (DSHB, M114, 1:500), DAPI (1:1000).

### Widefield microscopy

Images of immunostained ovaries mounted in SlowFade Gold antifade reagent (Invitrogen S36936) were acquired using a DeltaVision Elite microscope system (Applied Precision) equipped with a 100x oil immersion UPlanS-Apo objective (NA 1.4). Images were acquired as z-stacks with a step size of 0.5 μm. Fluorescence passed through a 435/48 nm; 525/48 nm; 597/45 nm; 632/34 nm band-pass filter for detection of respectively DAPI, Alexa Fluor 488, mCherry and Alexa Fluor 647 in sequential mode.

### Confocal microscopy

Images for Figure 4 were taken using an inverted Fluoview 1000 laser scanning microscope (Olympus) equipped with a 60× oil-immersion UPlanS-Apo objective (NA 1.2). The samples were excited at 404, 473, 559, and 635 nm, respectively, for DAPI and Alexa Fluor 488, 546, and 647. Light was guided to the sample via D405/473/559/635 dichroic mirror (Chroma). The pinhole was set at 115 μm. Fluorescence was passed sequentially through a 430–455-, 490–540-, 575–620-, 655–755-nm bandpass filter for detection of DAPI and Alexa Fluor 488, 546, and 647. Images were acquired as z-stacks with a step size of 0.5 μm

### Quantification

For each quantification one cell/germarium was considered. Images from a single cell (nucleus) were projected (max intensity) to capture all the centromeres present in the cell at a specific cell cycle phase. Image J software [56] was used to measure fluorescent intensity of CID in the following way: The background was subtracted from the projected image. Threshold was adjusted and the image. Size was adjusted, in order to eliminate unwanted objects. Following, the command “analyse particles” was used to select centromeres. Finally, integrated density (MGV*area) from each centromere foci were extracted and used as fluorescent intensity to measure the total amount of fluorescence per nucleus. Quantification of pMad and SEX-LETHAL positive cells was obtained counting the positive cells for each signal through the z-stack of each image. Statistical analysis was performed using prism software. Data distribution was assumed to be normal, but this was not formally tested. P value in each graph showed was calculated with unpaired t test with Welch’s correction.

## Acknowledgments

E.M.D. is funded by Science Foundation Ireland -PIYRA 13/YI/2187. B.L.C. is funded by Government of Ireland Postgraduate Fellowship 2018/1208 and by Science Foundation Ireland-PIYRA 13/YI/2187. A.A.D. was funded by Government of Ireland Postdoctoral Fellowship 2017/1324 and Science Foundation Ireland-PIYRA 13/YI/2187. The authors acknowledge the facilities and technical assistance of the Centre for Microscopy & Imaging at the National University of Ireland Galway (www.imaging.nuigalway.ie). Stocks were obtained from the Bloomington Drosophila Stock Center (NIH P40OD018537). Antibodies obtained from the Developmental Studies Hybridoma Bank, created by the NICHD of the NIH are maintained at The University of Iowa, Department of Biology, Iowa City, IA 52242. We thank Sylvia Erhardt for rabbit anti-CAL1 antibodies. We thank Annie Walshe for generation of the sheep anti-CENP-C aa 1-732 antibody.

## Notes

### Competing Interest Statement

The authors have declared no competing interest.

### Summary of Updates

Figure 5 and Figure S5 revised

## References

1. Horvitz HR, Herskowitz I. Mechanisms of asymmetric cell division: Two Bs or not two Bs, that is the question. Cell. Cell Press; 1992. pp. 237–255. doi:10.1016/0092-8674(92)90468-R

2. Knoblich JA. Mechanisms of Asymmetric Stem Cell Division. Cell. Cell Press; 2008. pp. 583–597. doi:10.1016/j.cell.2008.02.007

3. Knoblich JA. Asymmetric cell division: Recent developments and their implications for tumour biology. Nature Reviews Molecular Cell Biology. Nature Publishing Group; 2010. pp. 849–860. doi:10.1038/nrm3010

4. Clevers H. Stem cells, asymmetric division and cancer. Nature Genetics. Nature Publishing Group; 2005. pp. 1027–1028. doi:10.1038/ng1005-1027

5. Morrison SJ, Kimble J. Asymmetric and symmetric stem-cell divisions in development and cancer. Nature. 2006;441: 1068–1074. doi:10.1038/nature04956

6. Resende LPF, Jones DL. Local signaling within stem cell niches: insights from Drosophila. Curr Opin Cell Biol. 2012;24: 225–231. doi:10.1016/j.ceb.2012.01.004

7. Morrison SJ, Spradling AC. Stem cells and niches: mechanisms that promote stem cell maintenance throughout life. Cell. 2008;132: 598–611. doi:10.1016/j.cell.2008.01.038

8. Tran V, Lim C, Xie J, Chen X. Asymmetric division of Drosophila male germline stem cell shows asymmetric histone distribution. Science. 2012;338: 679–682. doi:10.1126/science.1226028

9. Wooten M, Snedeker J, Nizami ZF, Yang X, Ranjan R, Urban E, et al. Asymmetric histone inheritance via strand-specific incorporation and biased replication fork movement. Nat Struct Mol Biol. 2019;26: 732–743. doi:10.1038/s41594-019-0269-z

10. Xie J, Wooten M, Tran V, Chen BC, Pozmanter C, Simbolon C, et al. Histone H3 Threonine Phosphorylation Regulates Asymmetric Histone Inheritance in the Drosophila Male Germline. Cell. 2015;163: 920–933. doi:10.1016/j.cell.2015.10.002

11. Ma B, Trieu TJ, Cheng J, Zhou S, Tang Q, Xie J, et al. Differential Histone Distribution Patterns in Induced Asymmetrically Dividing Mouse Embryonic Stem Cells. Cell Rep. 2020;32. doi:10.1016/j.celrep.2020.108003

12. Lansdorp PM. Immortal Strands? Give Me a Break. Cell. 2007;129: 1244–1247. doi:10.1016/j.cell.2007.06.017

13. Musacchio A, Desai A. A molecular view of kinetochore assembly and function. Biology. Multidisciplinary Digital Publishing Institute; 2017. p. 5. doi:10.3390/biology6010005

14. Fukagawa T, Earnshaw WC. The Centromere: Chromatin foundation for the kinetochore machinery. Developmental Cell. Cell Press; 2014. pp. 496–508. doi:10.1016/j.devcel.2014.08.016

15. Allshire RC, Karpen GH. Epigenetic regulation of centromeric chromatin: Old dogs, new tricks? Nature Reviews Genetics. Nature Publishing Group; 2008. pp. 923–937. doi:10.1038/nrg2466

16. Mendiburo MJ, Padeken J, Fülöp S, Schepers A, Heun P. Drosophila CENH3 is sufficient for centromere formation. Science. 2011;334: 686–690. doi:10.1126/science.1206880

17. McKinley KL, Cheeseman IM. The molecular basis for centromere identity and function. Nature Reviews Molecular Cell Biology. Nature Publishing Group; 2016. pp. 16–29. doi:10.1038/nrm.2015.5

18. Mitra S, Srinivasan B, Jansen LET. Stable inheritance of CENP-A chromatin: Inner strength versus dynamic control. Journal of Cell Biology. Rockefeller University Press; 2020. doi:10.1083/JCB.202005099

19. Müller S, Almouzni G. Chromatin dynamics during the cell cycle at centromeres. Nature Reviews Genetics. Nature Publishing Group; 2017. pp. 192–208. doi:10.1038/nrg.2016.157

20. Dattoli AA, Carty BL, Kochendoerfer AM, Morgan C, Walshe AE, Dunleavy EM. Asymmetric assembly of centromeres epigenetically regulates stem cell fate. J Cell Biol. 2020;219. doi:10.1083/JCB.201910084

21. Ranjan R, Snedeker J, Chen X. Asymmetric Centromeres Differentially Coordinate with Mitotic Machinery to Ensure Biased Sister Chromatid Segregation in Germline Stem Cells. Cell Stem Cell. 2019;25: 666–681.e5. doi:10.1016/j.stem.2019.08.014

22. García del Arco A, Edgar BA, Erhardt S. In Vivo Analysis of Centromeric Proteins Reveals a Stem Cell-Specific Asymmetry and an Essential Role in Differentiated, Non-proliferating Cells. Cell Rep. 2018;22: 1982–1993. doi:10.1016/j.celrep.2018.01.079

23. Wooten M, Ranjan R, Chen X. Asymmetric Histone Inheritance in Asymmetrically Dividing Stem Cells. Trends Genet. 2019;36: 30–43. doi:10.1016/j.tig.2019.10.004

24. Carty BL, Dunleavy EM. Centromere assembly and non-random sister chromatid segregation in stem cells. Essays Biochem. 2020 [cited 23 Jul 2020]. doi:10.1042/ebc20190066

25. Erhardt S, Mellone BG, Betts CM, Zhang W, Karpen GH, Straight AF. Genome-wide analysis reveals a cell cycle-dependent mechanism controlling centromere propagation. J Cell Biol. 2008;183: 805–818. doi:10.1083/jcb.200806038

26. Goshima G, Wollman R, Goodwin SS, Zhang N, Scholey JM, Vale RD, et al. Genes required for mitotic spindle assembly in Drosophila S2 cells. Science. 2007;316: 417–421. doi:10.1126/science.1141314

27. Chen CC, Dechassa ML, Bettini E, Ledoux MB, Belisario C, Heun P, et al. CAL1 is the Drosophila CENP-A assembly factor. J Cell Biol. 2014;204: 313–329. doi:10.1083/jcb.201305036

28. Roure V, Medina-Pritchard B, Lazou V, Rago L, Anselm E, Venegas D, et al. Reconstituting Drosophila Centromere Identity in Human Cells. Cell Rep. 2019;29: 464–479.e5. doi:10.1016/j.celrep.2019.08.067

29. Medina-Pritchard B, Lazou V, Zou J, Byron O, Abad MA, Rappsilber J, et al. Structural basis for centromere maintenance by Drosophila CENP-A chaperone CAL 1. EMBO J. 2020;39: e103234. doi:10.15252/embj.2019103234

30. Schittenhelm RB, Althoff F, Heidmann S, Lehner CF. Detrimental incorporation of excess Cenp-A/Cid and Cenp-C into Drosophila centromeres is prevented by limiting amounts of the bridging factor Cal1. J Cell Sci. 2010;123: 3768–79. doi:10.1242/jcs.067934

31. Guo LY, Allu PK, Zandarashvili L, McKinley KL, Sekulic N, Dawicki-McKenna JM, et al. Centromeres are maintained by fastening CENP-A to DNA and directing an arginine anchor-dependent nucleosome transition. Nat Commun. 2017;8: 15775. doi:10.1038/ncomms15775

32. Falk SJ, Guo LY, Sekulic N, Smoak EM, Mani T, Logsdon GA, et al. CENP-C reshapes and stabilizes CENP-A nucleosomes at the centromere. Science. 2015;348: 699–703. doi:10.1126/science.1259308

33. Kirilly D, Xie T. The Drosophila ovary: An active stem cell community. Cell Research. Nature Publishing Group; 2007. pp. 15–25. doi:10.1038/sj.cr.7310123

34. Ables ET, Drummond-Barbosa D. Cyclin E controls Drosophila female germline stem cell maintenance independently of its role in proliferation by modulating responsiveness to niche signals. Development. Company of Biologists; 2013. pp. 530–540. doi:10.1242/dev.088583

35. Kao SH, Tseng CY, Wan CL, Su YH, Hsieh CC, Pi H, et al. Aging and insulin signaling differentially control normal and tumorous germline stem cells. Aging Cell. 2015;14: 25–34. doi:10.1111/acel.12288

36. Casanueva MO, Ferguson EL. Germline stem cell number in the Drosophila ovary is regulated by redundant mechanisms that control Dpp signaling. Development. 2004;131: 1881–1890. doi:10.1242/dev.01076

37. Rebecca Sheng X, Matunis E. Live imaging of the Drosophila spermatogonial stem cell niche reveals novel mechanisms regulating germline stem cell output. Development. 2011;138: 3367–3376. doi:10.1242/dev.065797

38. Salzmann V, Inaba M, Cheng J, Yamashita YM. Lineage tracing quantification reveals symmetric stem cell division in drosophila male germline stem cells. Cell Mol Bioeng. 2013;6: 441–448. doi:10.1007/s12195-013-0295-6

39. Song X, Wong MD, Kawase E, Xi R, Ding BC, McCarthy JJ, et al. Bmp signals from niche cells directly repress transcription of a differentiation-promoting gene, bag of marbles, in germline stem cells in the Drosophila ovary. Development. The Company of Biologists Ltd; 2004. pp. 1353–1364. doi:10.1242/dev.01026

40. Chau J, Kulnane LS, Salz HK. Sex-lethal Facilitates the Transition From Germline Stem Cell to Committed Daughter Cell in the Drosophila Ovary. 2009;132: 121–132. doi:10.1534/genetics.109.100693

41. Salz HK, Dawson EP, Heaney JD. Germ cell tumors: Insights from the Drosophila ovary and the mouse testis. Molecular Reproduction and Development. John Wiley and Sons Inc.; 2017. pp. 200–211. doi:10.1002/mrd.22779

42. Carroll CW, Milks KJ, Straight AF. Dual recognition of CENP-A nucleosomes is required for centromere assembly. J Cell Biol. 2010;189: 1143–1155. doi:10.1083/jcb.201001013

43. Hemmerich P, Weidtkamp-Peters S, Hoischen C, Schmiedeberg L, Erliandri I, Diekmann S. Dynamics of inner kinetochore assembly and maintenance in living cells. J Cell Biol. 2008;180: 1101–1114. doi:10.1083/jcb.200710052

44. Nechemia-Arbely Y, Miga KH, Shoshani O, Aslanian A, McMahon MA, Lee AY, et al. DNA replication acts as an error correction mechanism to maintain centromere identity by restricting CENP-A to centromeres. Nat Cell Biol. 2019;21: 743–754. doi:10.1038/s41556-019-0331-4

45. Zasadzińska E, Huang J, Bailey AO, Guo LY, Lee NS, Srivastava S, et al. Inheritance of CENP-A Nucleosomes during DNA Replication Requires HJURP. Dev Cell. 2018;47: 348–362.e7. doi:10.1016/j.devcel.2018.09.003

46. Pal S, Tyler JK. Epigenetics and aging. Science Advances. American Association for the Advancement of Science; 2016. doi:10.1126/sciadv.1600584

47. Sen P, Shah PP, Nativio R, Berger SL. Epigenetic Mechanisms of Longevity and Aging. Cell. Cell Press; 2016. pp. 822–839. doi:10.1016/j.cell.2016.07.050

48. Ermolaeva M, Neri F, Ori A, Rudolph KL. Cellular and epigenetic drivers of stem cell ageing. Nature Reviews Molecular Cell Biology. Nature Publishing Group; 2018. pp. 594–610. doi:10.1038/s41580-018-0020-3

49. Chen D, Kerr C. The Epigenetics of Stem Cell Aging Comes of Age. Trends in Cell Biology. Elsevier Ltd; 2019. pp. 563–568. doi:10.1016/j.tcb.2019.03.006

50. Nakagome Y, Abe T, Misawa S, Takeshita T, Iinuma K. The “loss” of centromeres from chromosomes of aged women. Am J Hum Genet. 1984;36: 398–404. Available: /pmc/articles/PMC1684436/?report=abstract

51. Yan D, Neumüller RA, Buckner M, Ayers K, Li H, Hu Y, et al. A regulatory network of Drosophila germline stem cell self-renewal. Dev Cell. 2014;28: 459–473. doi:10.1016/j.devcel.2014.01.020

52. Liu Y, Ge Q, Chan B, Liu H, Singh SR, Manley J, et al. Whole-animal genome-wide RNAi screen identifies networks regulating male germline stem cells in Drosophila. Nat Commun. 2016;7: 12149. doi:10.1038/ncomms12149

53. Neumüller RA, Richter C, Fischer A, Novatchkova M, Neumüller KG, Knoblich JA. Genome-wide analysis of self-renewal in Drosophila neural stem cells by transgenic RNAi. Cell Stem Cell. 2011;8: 580–593. doi:10.1016/j.stem.2011.02.022

54. Milagre I, Pereira C, Oliveira R, Jansen L. Reprogramming of Human Cells to Pluripotency Induces CENP-A Chromatin Depletion. Open Biol. 2020;10: 200227. doi:10.1101/2020.02.21.960252

55. Ambartsumyan G, Gill RK, Perez SD, Conway D, Vincent J, Dalal Y, et al. Centromere protein A dynamics in human pluripotent stem cell self-renewal, differentiation and DNA damage. Hum Mol Genet. 2010;19: 3970–3982. doi:10.1093/hmg/ddq312

56. Schneider CA, Rasband WS, Eliceiri KW. NIH Image to ImageJ: 25 years of image analysis. Nat Methods. 2012;9: 671–675. doi:10.1038/nmeth.2089

