## Supplemental Figures for "CENP-C regulates centromere assembly, asymmetry and epigenetic age in Drosophila germline stem cells"

**Supplementary Figure Legends: Carty et al.**

**Figure S1: Characterisation of CENP-C level in control *nanos-GAL4*, CENP-C RNAi, HA-CENP-C and HA-CENP-C; CENP-C RNAi lines.**

Immunofluorescent image of 5-day old (5d) $G_2$/prophase GSCs (circled) in (A-D) *nanos-GAL4* control, (A'-D') CENP-C RNAi and (A''-D'') HA-CENP-C; CENP-C RNAi stained with DAPI (cyan), CENP-C (green) and 1B1 (red). Scale bar = 10 μm. (E) Quantitation of total CENP-C fluorescent intensity (integrated density) per GSC in *nanos-GAL4*, CENP-C RNAi and HA-CENP-C; CENP-C RNAi (rescue). ***p<0.001. Scale bar = 10 μm. Error bars = SEM. (F-I) Immunofluorescence image of 5-day old (5d) $G_2$/prophase GSCs (circled) over-expressing HA-CENP-C stained with DAPI (cyan), CENP-C (green) and 1B1 (red). * denotes cap cells/GSC niche. GSCs are circled. Scale bar = 10 μm. (J) Quantitation of total CENP-C fluorescent intensity (integrated density) per GSC in HA-CENP-C. *p<0.05. Error bars = SEM.

**Figure S2: CENP-C is required for CAL1 localisation in GSCs, but it is not required for CID localisation at later stages of development.**

Immunofluorescent image of $G_2$/prophase GSCs (circled) in (A-C) *nanos-GAL4* and (A'-C') CENP-C RNAi stained for DAPI (cyan), 1B1 (red), CAL1 (yellow) and CENP-C (green). Centromeric CAL1 was identified as being colocalised with CENP-C. GSCs are circled. *denotes cap cells. Scale bar = 5 μm. (D) Quantitation of total centromeric CAL1 fluorescent intensity (integrated density) in *nanos-GAL4* and CENP-C RNAi. ***p<0.001. Error bars = SEM. (E-H) *bam-GAL4* and (E'-H') CENP-C RNAi stained with DAPI (cyan), CENP-C (yellow) and 1B1 (red). Circle marks region where knockdown begins. * denotes cap cells. Scale bar = 10 μm. (I-L) *bam-GAL4* and (I'-L') CENP-C RNAi stained with DAPI (cyan), CID (yellow) and 1B1 (red). Circle marks region where knockdown begins. * denotes cap cells. Scale bar = 10 μm.

**Figure S3: Quantitation of CID level in GSCs and CBs at S-phase.**

Quantitation of total CID fluorescent intensity (integrated density) in S-phase GSCs and CBs in *nanos-GAL4* and (A) CENP-C RNAi or (B) HA-CENP-C or (C) HA-CENP-C; CENP-C RNAi (rescue). Each point represents the total CID integrated density per GSC/CB nucleus. *p<0.05. **p<0.01. ns = non-significant. Error bars = SEM.

**Figure S4: Germaria with reduced CENP-C are not arrested in mitosis, but exhibit a higher frequency of S-phase cells.**

(A-D) *nanos-GAL4* and (E-H) CENP-C RNAi (severe GSC tumour phenotype) stained with DAPI (cyan), VASA (grey) and H3PS10 (red). * denotes cap cells. Scale bar = 10 μm. (I-L) *nanos-GAL4* and

(M-P) CENP-C RNAi stained with DAPI (cyan), EdU (yellow) and 1B1 (red). * denotes cap cells. White dashed lines outline two EdU positive cysts. Scale bar = 10 μm. (Q) Quantitation of the number of EdU positive cysts per germarium. Graph represents the median ± Standard Deviation. One positive hit was quantified as a single EdU positive GSC/CB, 2-cell cysts (2cc), 4-cell cysts (4cc) or 8-cell cysts (8cc). ***p<0.001.  Error bars = Standard Deviation.

**Figure S5: A method of measuring female GSC self-renewal versus differentiation.**

(A-C) *Wild type* (*OregonR*) (5-, 10- and 20-days old) and (D) *wild type* (#36303, RNAi isogenic control) germaria stained with DAPI (cyan), SXL (red) and pMad (yellow).  *denotes cap cells. Scale bar = 10 μm. White dashed circles highlight SXL or pMad positive cells. (E) Quantitation of the number of pMad positive (left, yellow) and SXL positive (right, red) per germarium. (F) Ratio of the number of SXL:pMad positive cells per germarium. ***p<0.001, *p<0.05. Error bars = SEM.

### Figure S1

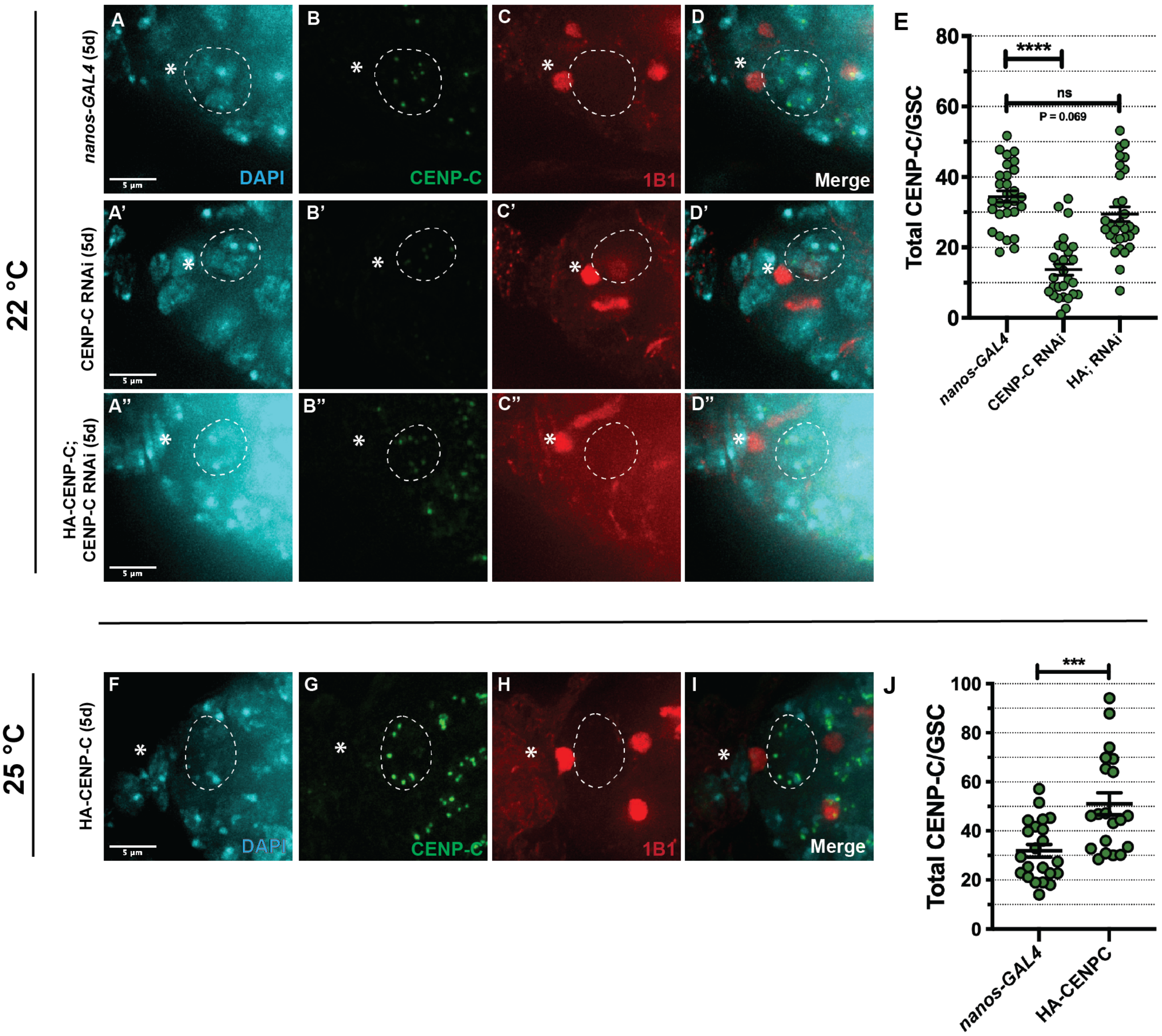

22 °C
25 °C
nanos-GAL4 (5d)
CENP-C RNAi (5d)
HA-CENP-C; CENP-C RNAi (5d)
HA-CENP-C (5d)
A
B
C
D
A'
B'
C'
D'
A"
B"
C"
D"
F
G
H
I
DAPI
CENP-C
1B1
Merge
5 µm
E
Total CENP-C/GSC
****
ns
P = 0.069
nanos-GAL4
CENP-C RNAi
HA; RNAi
J
***
HA-CENPC

### Figure S2

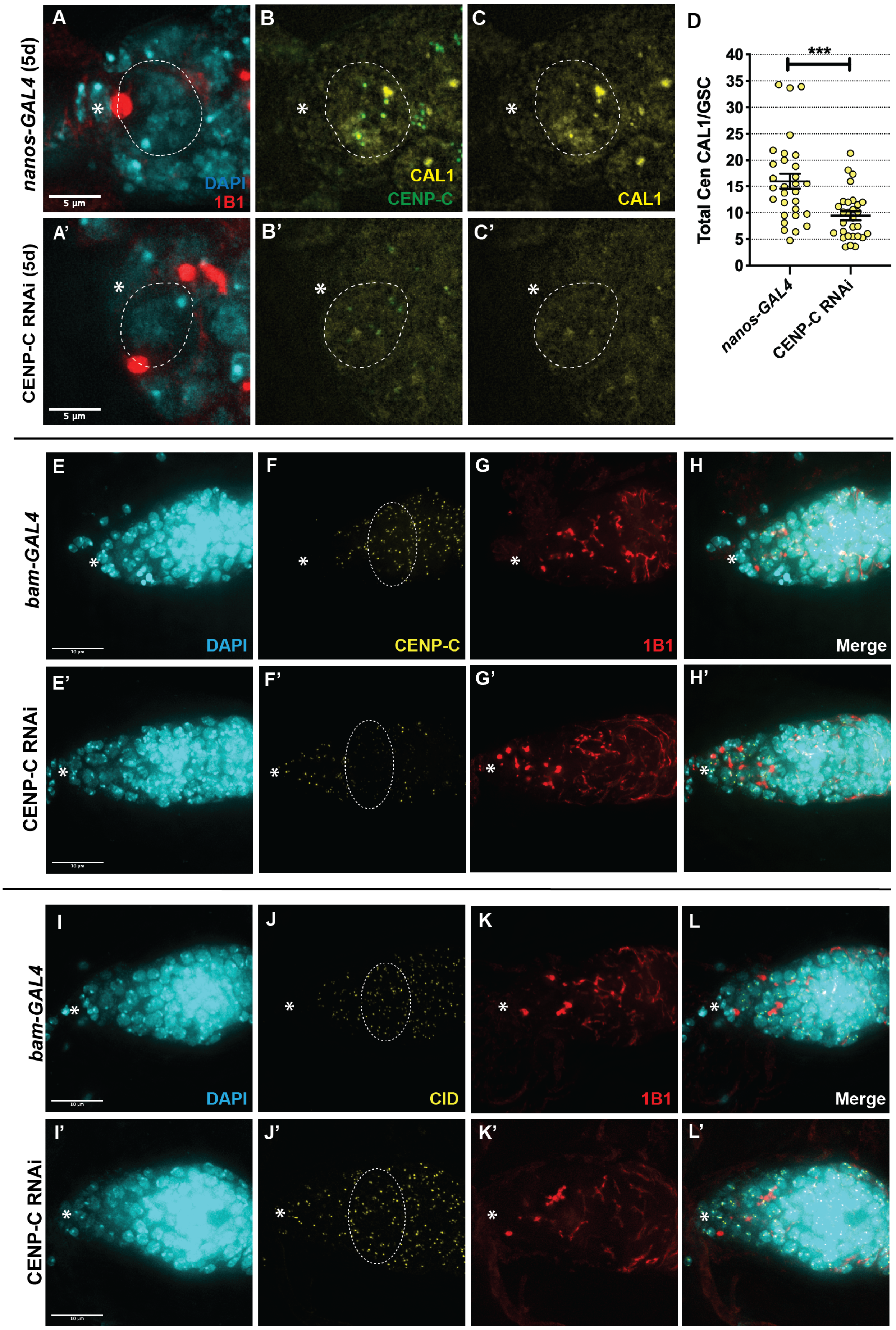
A
B
C
D
nanos-GAL4 (5d)
DAPI
1B1
5 µm
CAL1
CENP-C
CAL1
A'
B'
C'
CENP-C RNAi (5d)
5 µm
***
Total Cen CAL1/GSC
40
35
30
25
20
15
10
5
0
nanos-GAL4
CENP-C RNAi
E
F
G
H
bam-GAL4
DAPI
CENP-C
1B1
Merge
E'
F'
G'
H'
CENP-C RNAi
I
J
K
L
bam-GAL4
DAPI
CID
1B1
Merge
I'
J'
K'
L'
CENP-C RNAi

### Figure S3

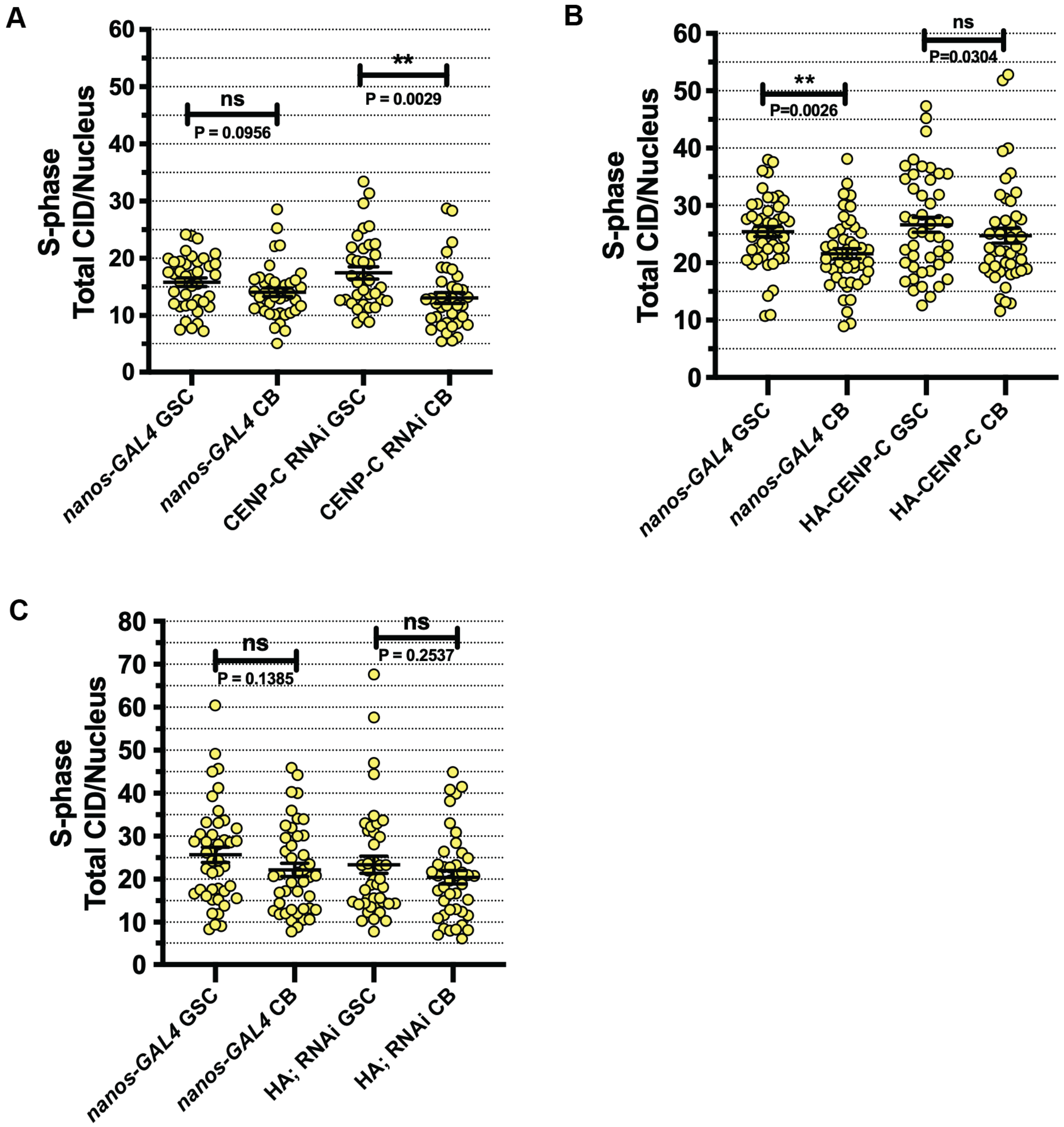
A
S-phase Total CID/Nucleus
60
50
40
30
20
10
0
ns
P = 0.0956
**
P = 0.0029
nanos-GAL4 GSC
nanos-GAL4 CB
CENP-C RNAi GSC
CENP-C RNAi CB
B
S-phase Total CID/Nucleus
**
P=0.0026
ns
P=0.0304
nanos-GAL4 GSC
nanos-GAL4 CB
HA-CENP-C GSC
HA-CENP-C CB
C
S-phase Total CID/Nucleus
80
70
ns
P = 0.1385
ns
P = 0.2537
nanos-GAL4 GSC
nanos-GAL4 CB
HA; RNAi GSC
HA; RNAi CB

### Figure S4

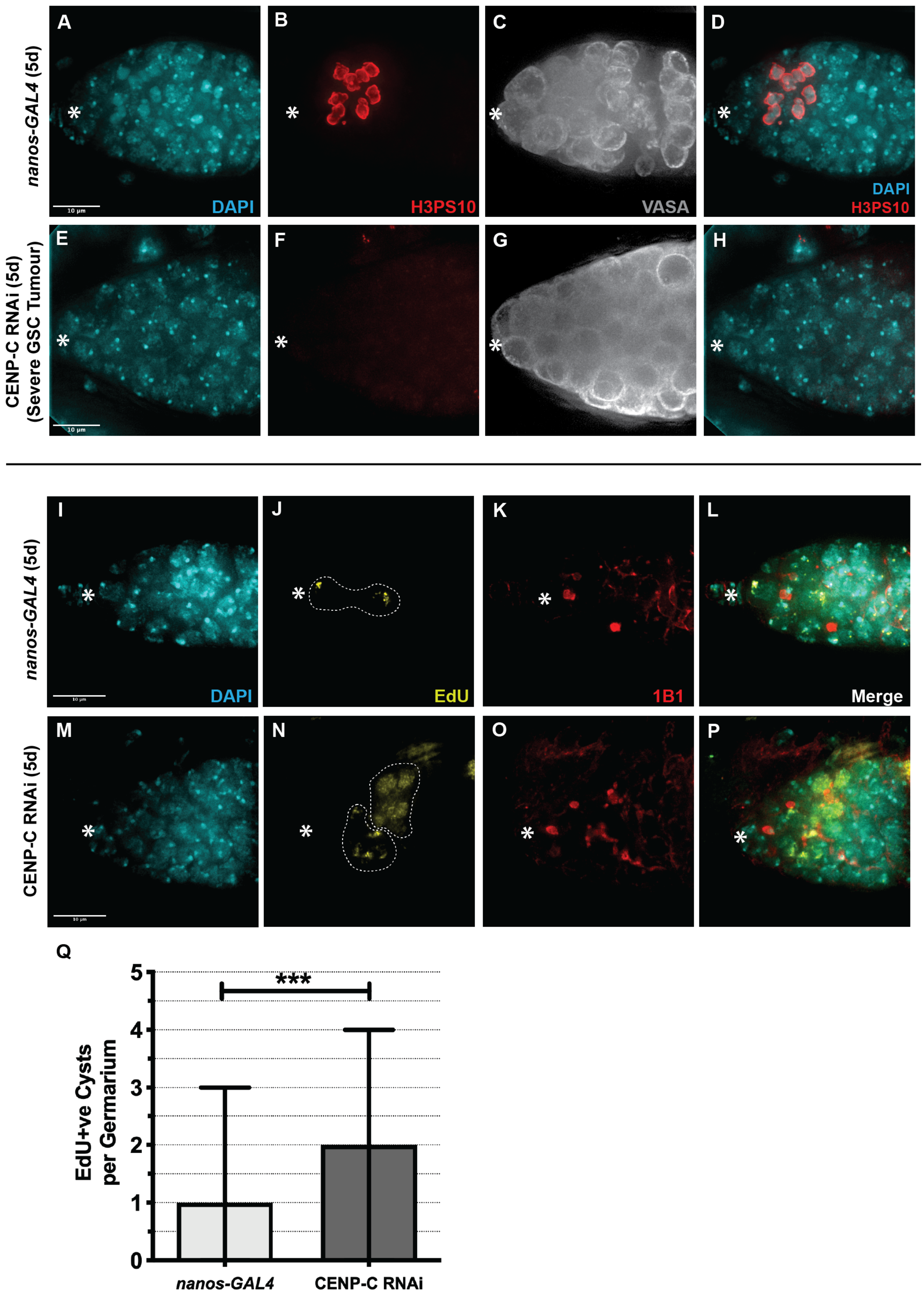

nanos-GAL4 (5d)
CENP-C RNAi (5d) (Severe GSC Tumour)
A
B
C
D
DAPI
H3PS10
VASA
DAPI
H3PS10
E
F
G
H
10 μm
I
J
K
L
DAPI
EdU
1B1
Merge
nanos-GAL4 (5d)
M
N
O
P
CENP-C RNAi (5d)
Q
EdU+ve Cysts per Germarium
0
1
2
3
4
5
***
nanos-GAL4
CENP-C RNAi

### Figure S5

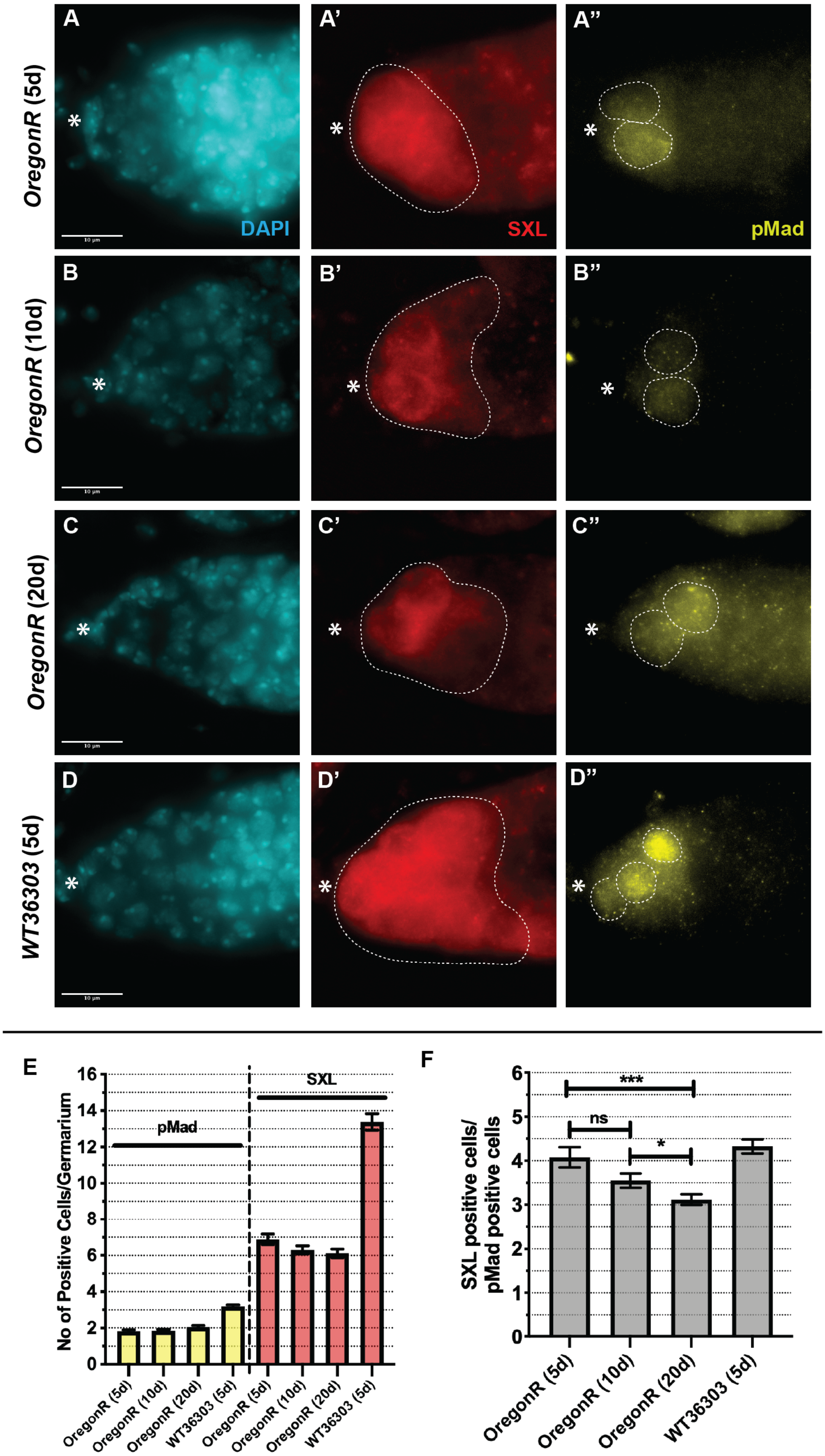
A
A'
A''
B
B'
B''
C
C'
C''
D
D'
D''
OregonR (5d)
OregonR (10d)
OregonR (20d)
WT36303 (5d)
DAPI
SXL
pMad
E
No of Positive Cells/Germarium
pMad
SXL
OregonR (5d)
OregonR (10d)
OregonR (20d)
WT36303 (5d)
F
SXL positive cells/
pMad positive cells
ns
*
***
